# Non-invasive *Bdnf* mRNA therapy improves cognition in ageing and Alzheimer’s mouse models

**DOI:** 10.64898/2026.04.19.719519

**Authors:** Maria Bergamasco, Teleri Clark, Lipin Loo, Kazuma Fujikake, Rebecca Carr, Hannah Scarborough, Anna Ponta, R.M. Damian Holsinger, Greg Neely

## Abstract

Messenger RNA (mRNA) therapeutics have rapidly emerged as a transformative approach for treating a range of health challenges. Accelerated by the success of mRNA-lipid nanoparticle (LNP) vaccines during the COVID-19 pandemic, this platform holds promise beyond immunisation for the transient expression of therapeutic proteins in targeted tissues. Despite this promise, non-invasive delivery of mRNA to the brain, as with most therapeutics, remains a challenge due to the impermeability of the blood brain barrier. Here, we present a novel strategy to deliver neurotrophic factors to the brain via intranasal delivery of mRNA-LNP. As a proof of concept, we demonstrate that intranasal delivery of mRNA encoding the neurogenic factor BDNF (Brain Derived Neurotrophic Factor) enhances memory performance in both aged mice and a transgenic mouse model of Alzheimer’s disease. This approach offers a promising platform for delivering therapeutic proteins to the brain and opens new avenues for treating age-related and neurodegenerative disorders.

## INTRODUCTION

Messenger RNA (mRNA) therapeutics have been propelled into the global spotlight by the rapid development and deployment of mRNA-based vaccines during the COVID-19 pandemic. These vaccines not only demonstrated the feasibility of large-scale mRNA production and delivery but also validated lipid nanoparticles (LNPs) as a clinically viable platform for nucleic acid transport. Beyond infectious diseases, the modularity, safety and rapid manufacturability of mRNA-LNP systems have catalysed interest in their application across a spectrum of conditions, including cancer, metabolic disorders and neurological diseases, with over 70 mRNA-based therapies currently in clinical trial (Reviewed in ^[1]^)

Age-related cognitive decline and neurodegenerative disorders such as Alzheimer’s disease (AD), represent some of the most pressing challenges in medicine. Notably, mild cognitive impairment in midlife (age 50-60) predicts severe decline in later life ^[2]^. As such, interventions that delay or mitigate cognitive deterioration have the potential to significantly reduce the societal and economic burdens of an ageing population. Current therapeutic strategies to manage age-related cognitive decline and neurodegenerative disorders, however, largely focus on symptomatic treatment or are only modestly disease modifying.

One major obstacle to improved therapeutics for these conditions is drug delivery to the brain, constrained by the blood-brain-barrier (BBB). To circumvent this barrier, several direct delivery methods have been developed, including intravenous-guided intracerebral injection, intrathecal administration and convection-enhanced delivery. While these approaches enable precise targeting to specific brain regions, they are inherently invasive, often requiring surgical procedures and posing risks such as infection, tissue damage and limited patient compliance. In contrast, systemic delivery methods, such as intravenous or subcutaneous injection, are less invasive and more amenable to repeated administration. These alternative approaches, however, suffer from poor brain bioavailability and their systemic distribution increases the likelihood of off-target effects in peripheral organs. Intranasal delivery represents a promising non-invasive alternative that bypasses the BBB by exploiting the olfactory and trigeminal neural pathways. This route enables direct access to the brain from the nasal cavity, facilitating rapid transport of molecules to the central nervous system without systemic circulation. Recent studies have demonstrated its utility in delivering peptides^[3]^, nanoparticles^[4]^ and nucleic acids^[5]^ to the brain.

Neurotrophic factors, such as brain-derived growth factor (BDNF), nerve growth factor (NGF) and glial cell line-derived neurotrophic factor (GDNF) are essential regulators of neuronal survival, synaptic plasticity and cognitive function. Among these, BDNF is a particularly potent modulator of learning and memory, shown to enhance synaptic density and neurogenesis^[6]^. Reduced *BDNF* expression is observed in the brains of individuals with AD and other neurodegenerative conditions, compared to healthy controls^[7]^. Consistently, elevating BDNF levels has been found to improve cognition in AD animal models^[8]^ and higher baseline serum BDNF levels are associated with a reduced risk of developing dementia^[9]^. Furthermore, in an AD clinical trial, cerebrolysin plus donepezil was shown to increase serum BDNF, with larger BDNF increases correlated with greater cognitive improvements^[10]^.

Here, we report the development of an intranasally delivered mRNA-LNP, designed to bypass systemic circulation and directly target the central nervous system. As a proof of concept, we use this platform to deliver mRNA encoding *Bdnf* to the brains of aged mice and 5xFAD AD animals. Our findings provide preclinical evidence supporting the feasibility and efficacy of this strategy in mitigating cognitive decline, offering a promising avenue for the treatment of age-related and neurodegenerative cognitive disorders.

## RESULTS

### Intranasal administration of LNP-mRNA can target the brain

Therapeutic delivery to the CNS is constrained by the blood-brain-barrier (BBB). Olfactory neurons project axons through the cribriform plate to form a direct interface between olfactory stimuli in the environment and the CNS. To exploit this feature, we used intranasal delivery of the clinically validated BioNTech COVID-19 mRNA backbone to deliver mRNA to the brain^[11]^. *In vitro* transcribed mRNA was encapsulated into LNPs and administered intranasally to mice under light anaesthesia **(Figure. 1A-B)**. To quantify the delivery efficiency and tissue distribution of delivered LNPs and mRNA, mice were administered with mRNA encoding a luciferase reporter and the near infrared dye, DiR, was incorporated into the LNP formulation. IVIS imaging of the skull, brain and peripheral organs of treated mice showed Luciferase expression in the respiratory epithelium **(Figure. 1C)**, with DiR detected throughout the respiratory and olfactory epithelia **(Figure. 1D)**. Importantly, no signal was detected in peripheral organs, including the lungs. This indicates that the LNP-mRNA did not migrate beyond the nasal cavity or enter the lower respiratory tract, a potential concern if material were to drain posteriorly into the throat. No signal was detected in any tissue in control animals.

**Figure 1:**
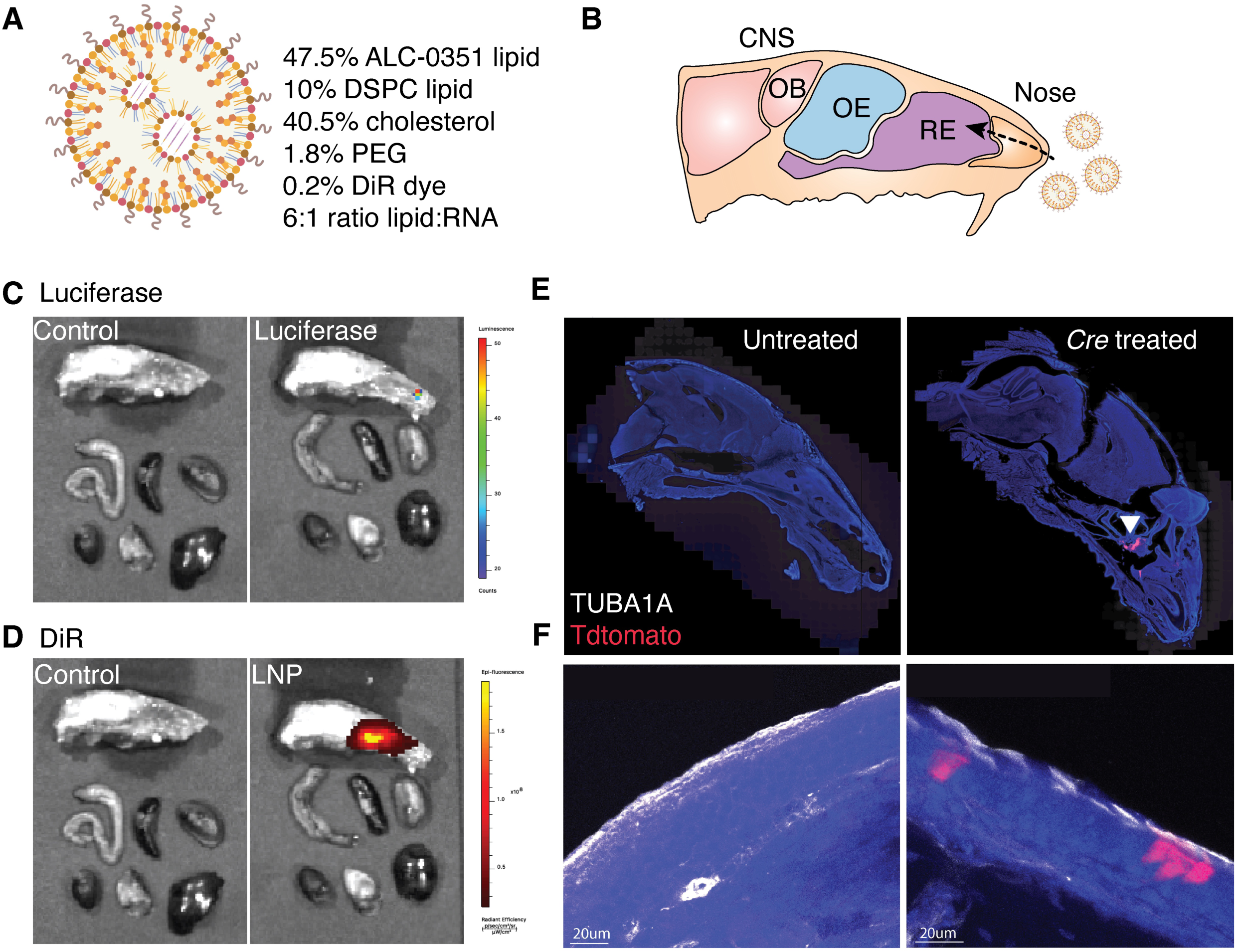
Intranasal delivery of LNP-mRNA to target the brain. **A.** Illustration of LNP formulation **B.** Diagram of sagittal view of the mouse skull, showing LNP delivery via the nose to the central nervous system (CNS). Respiratory epithelium (RE), olfactory epithelium (OE) and olfactory bulb (OB). **C-D.** IVIS imaging for Luciferase (C) and DiR dye (D) expression in the skull and peripheral organs of mice treated with control or luciferase mRNA-containing LNP. **E-F.** Confocal images of sagittal sections of the skull in untreated controls or animals treated with LNP-mRNA encoding *Cre recombinase* (E). Zoomed in images showing Tdtomato expression (red) and TUBA1A endothelial cell markers (white) (F). Scale bar = 200 μm.

To assess whether delivered mRNA cargo was functionally active *in vivo* we used *tdTomato* reporter mice carrying a floxed STOP cassette upstream of the *tdTomato* gene ^[12]^. Intranasal administration of LNP-mRNA encoding Cre recombinase gave robust *tdTomato* expression within epithelial cells (marked by α tubulin) within the olfactory and respiratory epithelia **(Figure. 1E,F)**. No signal was detected in untreated control mice. This recombination-dependent activation confirmed successful mRNA translation and enzymatic activity in situ, demonstrating that intranasally delivered mRNA-LNPs can drive functional protein expression.

### *Bdnf* mRNA promotes neural stem cell proliferation and neuronal differentiation

BDNF is a neurotrophin with potent neurorestorative and synaptic plasticity-enhancing effects. Despite strong evidence in the literature for the benefits of elevated BDNF levels across neurodegenerative disorders, its efficacy in clinical settings has been limited by its inability to cross the BBB. Given its promise as a therapeutic, we selected *Bdnf* mRNA for proof-of-concept assessments of our LNP-mRNA platform.

Endogenous BDNF is synthesised as a precursor (pre-pro-BDNF) that undergoes sequential cleavage to generate either pro-BDNF or mature BDNF (mBDNF). Importantly, the ratio of mBDNF to pro-BDNF influences cognitive function, with excess pro-BDNF associated with synaptic dysfunction and impaired learning and memory^[13]^. To avoid these deleterious effects and increase only mature BDNF, we engineered an mRNA construct encoding the mature BDNF domain fused to its native signal peptide **(Figure. 2A)**.

**Figure 2:**
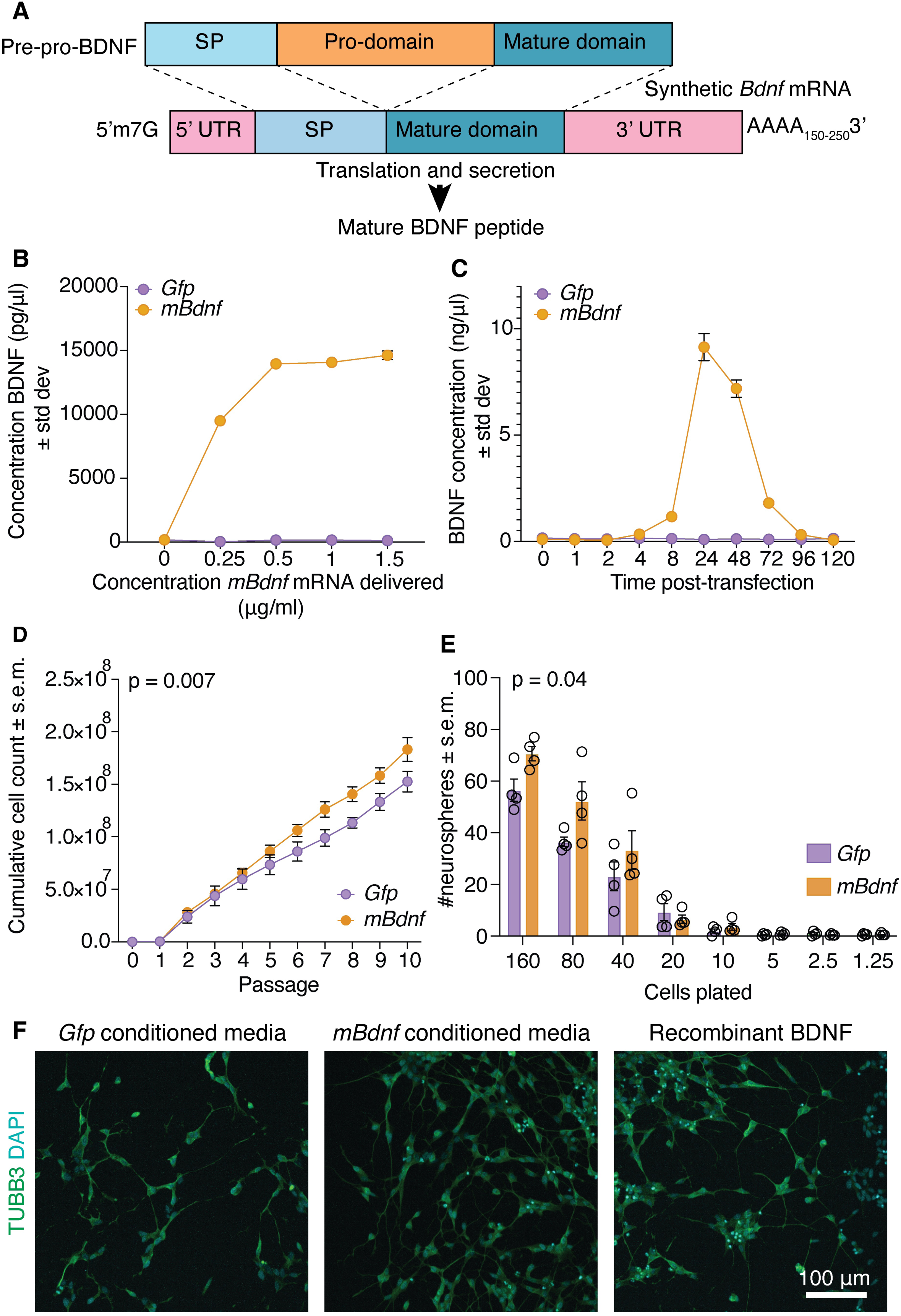
*Bdnf* mRNA dynamics and effect on NSPCs and SH-SY5Y cells. **A.** Schematic of synthetic *Bdnf* mRNA. The endogenous BDNF signal peptide (SP) was directly fused to the mature domain and bordered by 5’ and 3’ UTR sequences. A 5’m7G cap and polyA tail promote mRNA stability. **B.** ELISA analysis of the concentration of BDNF (pg/μl) in the media of HEK293T cells transfected with *Gfp* or *mBdnf* mRNA. Media collected 24 hr after transfection. Average of 2 technical replicates. **C.** ELISA analysis of the concentration of BDNF (pg/μl) in the media of HEK293T cells transfected with *Gfp* or *mBdnf* mRNA. Media collected at various timepoints after transfection and replenished. Average of two technical replicates. **D.** Cumulative cell counts of adult hippocampal NSPCs cultured in conditioned media from HEK293T cells transfected with *Gfp* or *mBdnf* mRNA. Data presented as mean ± s.e.m. N = 4x NSPC cultures derived from 4x adult mouse hippocampi. Data analysed using a two-way ANOVA with Sidak post-hoc correction. **E.** Number of secondary neurospheres derived from adult hippocampal NSPCs cultured in conditioned media from HEK293T cells transfected with *Gfp* or *mBdnf* mRNA plates, at a dilution range. Data presented as mean ± s.e.m. N = NSPCs from 4x animals per mRNA group. Data analysed using a two-way ANOVA with Sidak post-hoc correction. **F.** TUBB3 (green) and DAPI (blue) staining of SH-SY5Y-derived neurons treated with recombinant BDNF or conditioned media from *Gfp* or *mBdnf* mRNA-treated HEK293T cells. Scale bar = 100 μm.

To confirm that this modified mRNA was secreted and produced detectable protein, we performed an ELISA assay on cell culture supernatant from HEK293T cells transfected with *mBdnf* mRNA or *Gfp* control mRNA. Substantial amounts of BDNF were detected in the media of cells transfected with *mBdnf* mRNA at a range of doses. No BDNF was detected in the media of *Gfp* transfected cells **(Figure. 2B)**. A time course of transfection showed that BDNF secretion was detectable from 2 hr post-transfection, peaked at 24 hr post-transfection and remained detectable in cell culture media for up to 4 days post-transfection **(Figure. 2C)**, indicating sustained protein production from the delivered mRNA.

Given that mBDNF is an established regulator of neural stem cell (NSC) proliferation^[14]^ and neuronal differentiation^[15]^, we assessed whether our modified BDNF retained biological activity by examining its effects on adult hippocampal neural stem and progenitor cells (NSPCs) and SH-SY5Y neuroblastoma cells. NSPCs were grown in conditioned media from HEK293T cells transfected with *mBdnf* or *Gfp* mRNA. Compared to *Gfp* controls, *mBdnf* treated cells showed enhanced proliferation over consecutive passages (p = 0.007; **Figure. 2D**) and gave rise to more secondary neurospheres across plating dilutions (p = 0.04; **Figure. 2E**), indicating enhanced self-renewal.

To assess neuronal differentiation, SH-SY5Y cells were induced to differentiate in the presence of either recombinant BDNF or conditioned media from *mBdnf* or *Gfp*-transfected HEK293T cells. Compared to cells that did not receive BDNF (*Gfp* controls), recombinant BDNF (50 ng/ml) and *mBdnf* conditioned media-treated cells showed more pronounced neurite outgrowth and connectivity **(Figure. 2F)**, indicating that *mBdnf* mRNA results in the production of bioactive BDNF that can drive neuronal differentiation similarly to the recombinant protein.

### *Bdnf* mRNA treatment in aged mice improves memory and cognition

Having confirmed *in vitro* that our mRNA construct produced functional BDNF, we next evaluated its effects *in vivo. mBdnf* and *Gfp* control mRNA was encapsulated into LNPs (92 and 76 nm in diameter respectively, based on a zetasizer) **(Figure. S1A)**. 24 hrs after intranasal delivery, RTqPCR was used to assess *mBdnf* mRNA levels in treated mice. Primers were designed that spanned the novel fusion site between the BDNF signal peptide and mature domain to distinguish delivered from endogenous *Bdnf* mRNA. Remarkably, *mBdnf* transcript levels were 1000-50,000-fold elevated in the respiratory epithelium, olfactory epithelium and olfactory bulb of *mBdnf*-treated vs *Gfp* control mice (p <0.0001 – 0.04; **Figure. S1B**), demonstrating very large increases in *mBdnf* expression and verifying delivery to the CNS. Similarly, BDNF levels were significantly higher in *mBdnf*-treated brains, compared to *Gfp*-treated controls, by both western immunoblotting (**Figure. S1C,D**, p =0.045) and immunohistochemistry in the hippocampus **(Figure. S1E)**.

We next assessed the functional effect of *mBdnf* treatment on behaviour. As in humans, aged mice display cognitive deficits in behavioural assessments of learning and memory^[16]^, which correlates with an age-related decline in brain BDNF levels^[17]^. Aged (18 mn) mice were treated with intranasal drops of *Gfp* or *mBdnf* LNP-mRNA and underwent behavioural assessments of cognition; the open field test, novel object recognition test, Y maze for working memory and Y maze for spatial memory **(Figure. 3A)**. All tests were also performed on untreated young (3mn) mice.

**Figure 3:**
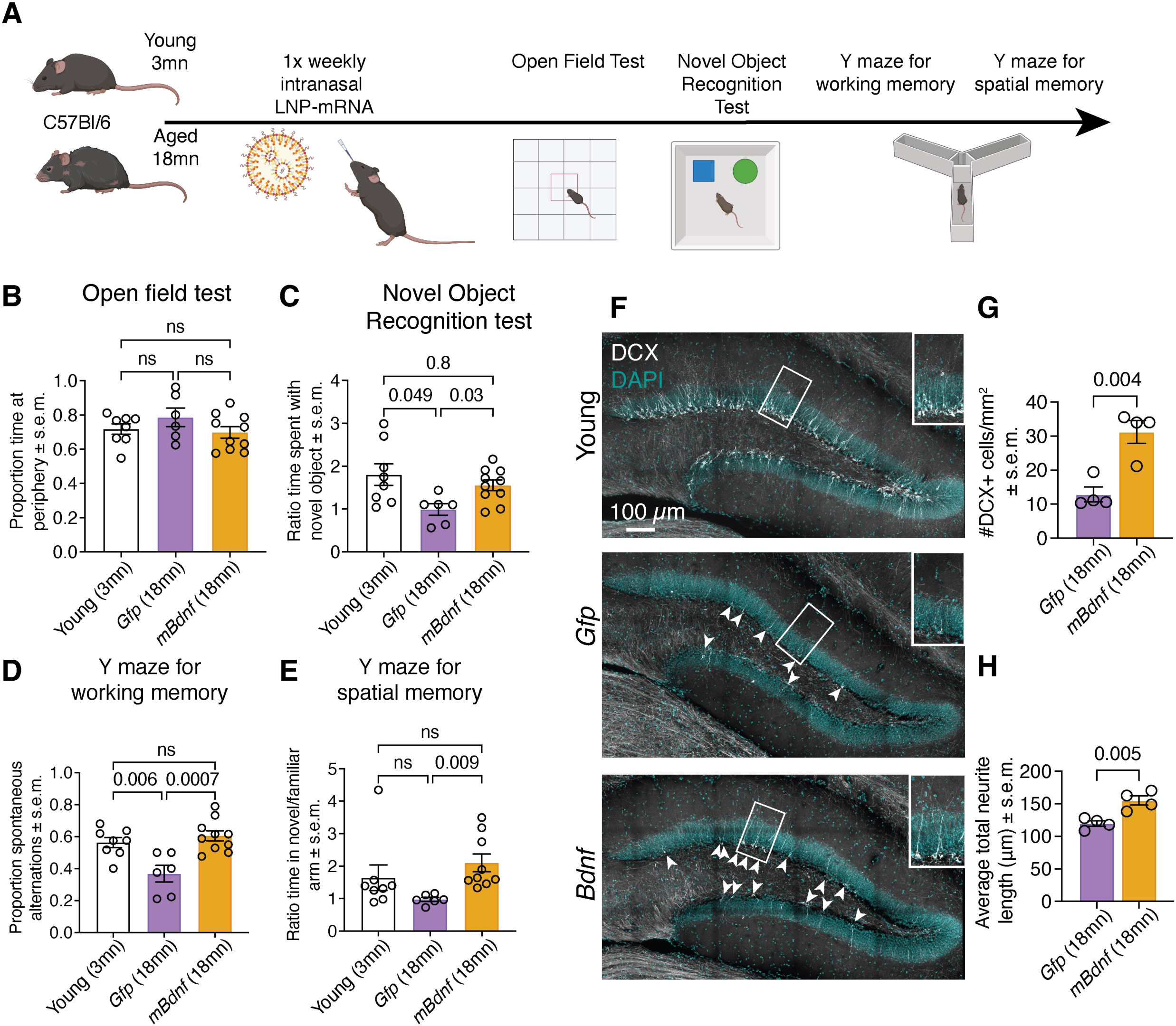
*Bdnf* mRNA-LNP treatment in aged mice improves memory performance. **A.** Schematic of the treatment schedule of LNP-mRNA in aged (18 mn mice). Animals received 1x weekly dose of 1 μg mRNA-LNP 24 hr prior to a behavioural test: open field test, novel object recognition, Y maze for working memory and Y maze for spatial memory. **B-E.** Analysis of behavioural tests in young (3mn) mice or aged (18 mn) mice treated with mRNA encoding *Gfp* or *mBdnf*. Proportion of time spent at the periphery of the arena in the open field (A), ratio of time spend with the novel/familiar object in the novel object recognition test (C), proportion of spontaneous alternations in the Y maze for working memory (D) and ratio of time spent in the novel/familiar arm in the Y maze for spatial memory (E) were assessed. In all graphs each circle represents an individual animal. Data presented as mean ± s.e.m. N = 6-10 animals per age and treatment group. Data analysed using a one-way ANOVA with Dunnett post hoc correction. **F**. Doublecortex (DCX) (white) and DAPI (blue) staining (F) in hippocampal sections from young (3mn) mice or aged (18mn) mice treated with LNP-mRNA encoding *Gfp* or m*Bdnf.* **G,H** Number per mm^2^ (G) and average neurite total length (μm) (H) of DCX+ cells in *Gfp* and *mBdnf*-treated 18mn mouse hippocampal sections. Each circle in B-E, G and H represents an individual animal (B-E) or hippocampal sections from an individual animal (G,H). N = 4-10 samples per group. Data analysed using a one-way ANOVA with Dunnett post-hoc correction (B-E) or unpaired t-test (G,H).

In the open field test, there was no effect across age or treatment on the proportion of time spent at the periphery of the arena (p = 0.2-0.7; **Figure. 3B**). Both *Gfp* and *mBdnf-*treated aged mice travelled a shorter total distance and had a lower average speed than young mice (p = 0.0006-0.09; **Figure. S2A,B**). In the novel object recognition test, *Gfp*-treated aged animals spent less time with the novel object compared to young animals (p = 0.049), while *mBdnf* treated mice behaved similarly to young mice (p = 0.8) and performed significantly better than aged *Gfp*-treated mice (p = 0.03; **Figure. 3C**). Similarly in the Y maze for working memory, *Gfp*-treated aged mice performed poorly compared to both young mice (p = 0.006) and *mBdnf-*treated aged counterparts (p = 0.0007). By comparison *mBdnf* treated aged mice performed similarly to young animals (p = 0.7; **Figure. 3D**). In the Y maze for spatial memory *mBdnf* treated animals also performed better than *Gfp-*treated aged controls (p = 0.009) and again performed similarly to young mice (p = 0.7; **Figure. 3E**). These results demonstrate that a once-a-week dose of intranasal *mBdnf* mRNA-LNP therapy can significantly improve age-related cognitive decline.

Doublecortin (DCX) is a marker for neurogenesis^[18]^, expressed in newly generated neurons and reflective of plasticity and learning. Further reflective of improved memory performance, *mBdnf*-treated aged mice showed a 2.4-fold increase in the number of DCX positive cells in the hippocampus compared to *Gfp*-treated controls (p = 0.004; **Figure. 3F,G**). The DCX+ cells of *mBdnf*-treated aged animals were also more arborised than in *Gfp* treated controls (p = 0.005; **Figure. 3H**). Notably the number and arborisation of DCX+ cells in both aged treatment groups was less than in young mice. This indicates that following *mBdnf* mRNA-LNP therapy, neurogenesis is not completely restored to that of a young animal, however the effects of intranasal *mBdnf* mRNA-LNP are none-the-less sufficient to return cognitive performance to a youthful state.

Notably, mice treated with *mBdnf* mRNA-LNP therapy appeared healthy and showed no major side effects. Importantly, our LNP therapy contains PEG, and anti-PEG antibodies have been observed in patients treated with other PEGylated drugs^[19]^. Since *mBdnf* mRNA-LNP therapy requires repeated intranasal dosing, development of anti-PEG antibodies may impair therapeutic efficacy over time^[20]^. To address this concern, we assessed the serum of LNP-treated and untreated animals by ELISA; anti-PEG antibodies were not detected in the sera of either group **(Figure. S2C)**.

### *Bdnf* mRNA treatment in 5xFAD mice improves memory and cognition

BDNF mRNA and protein levels are reduced in postmortem AD brains and increasing BDNF levels in AD animal models and in human AD patients improve cognitive performance. To test *mBdnf* LNP-mRNA as a potential therapeutic for AD, we used the 5xFAD mouse model^[21]^. 5xFAD mice serve as a rapid onset “humanised” model of AD, with AD-associated brain pathologies detectable from 2 mn of age and cognitive decline apparent from 4 mn^[21–22]^.

To evaluate early intervention, 5xFAD animals were treated with *Gfp* or *mBdnf* mRNA-LNP twice weekly from 2 mn of age. From 4 mn of age animals underwent behavioural assessments, continuing LNP-mRNA doses during testing **(Figure. 4A)**. No significant difference was observed between treatment groups in the proportion of time spent at the periphery of the open field test (p = 0.2; **Figure. 4B**), however, across the novel object recognition test, Y maze for working memory and Y maze for spatial memory, *mBdnf*-treated animals performed significantly better than *Gfp* treated counterparts (p = 0.002-0.02; **Figure. 4C-E**). No difference was observed between treatment groups in the average speed or total distance travelled in any test **(Figure. S3A-D)**. As in aged mice, no anti-PEG antibodies were found in the serum of either treated group **(Figure. S3E)**.

**Figure 4:**
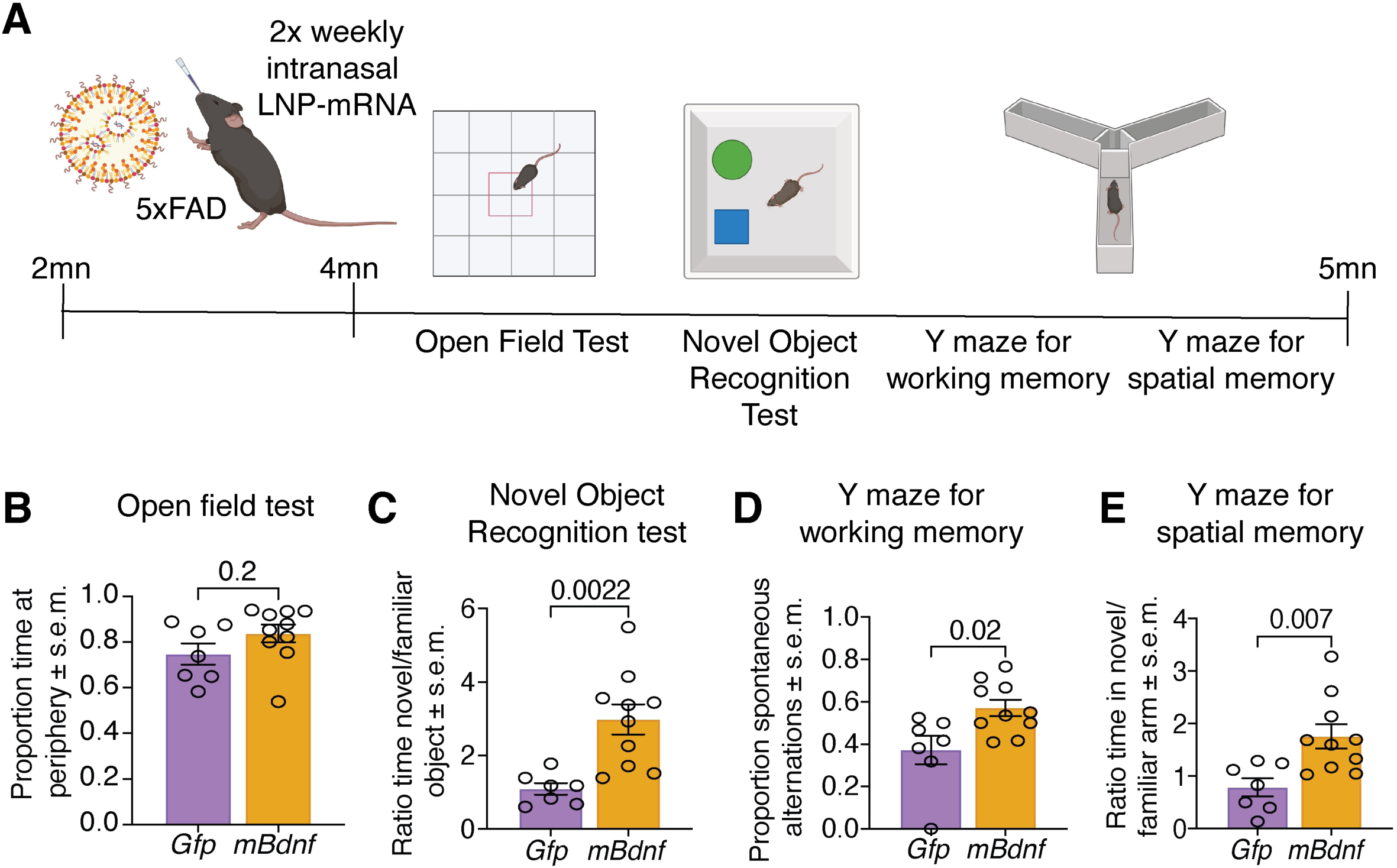
*Bdnf* mRNA-LNP treatment in 5xFAD mice improves memory performance. **A.** Schematic of the treatment schedule of LNP-mRNA in 5xFAD mice. Animals were treated 2x weekly from 2-4 mn of age. From 4 mn of age animals began behavioural assessments and continued LNP-mRNA treatment throughout behavioural testing. Animals underwent one behavioural test 24 hr after an LNP-mRNA dose. **B-E.** Analysis of behavioural tests in 5xFAD mice treated with mRNA encoding *eGfp* or *Bdnf*. Proportion of time spent at the periphery of the arena in the open field (B), ratio of time spend with the novel/familiar object in the novel object recognition test (C), proportion of spontaneous alternations in the Y maze for working memory (D) and ratio of time spent in the novel/familiar arm in the Y maze for spatial memory (E) were assessed. In all graphs each circle represents an individual animal. Data presented as mean ± s.e.m. N = 6-10 animals per age and treatment group. Data analysed using a one-way ANOVA with Dunnett post hoc correction.

Animals also underwent a histological assessment of the hippocampus. The role of amyloid plaques in AD is controversial, however therapeutic antibodies targeting plaques do show some clinical efficacy and are approved for clinical use (Reviewed in ^[23]^). We observed no difference in the amyloid plaque load (p = 0.3) or number of GFAP+ cells (p = 0.6; **Figure. 5A-C**) between treatment groups, indicating that *mBdnf* treatment did not impact this aspect of disease progression. DCX staining, however, showed that *mBdnf-*treated 5xFAD mice had more DCX+ cells within the hippocampus (p = 0.004) with a greater average total neurite length (p= 0.04; **Figure. 5D-F**). Similarly, *mBdnf*-treated animals had more NeuN+ cells in the CA3 and dentate gyrus regions of the hippocampus compared to *Gfp*-treated controls (p = 0.004 and 0.008; **Figure. 5G-L**). Overall, this suggests that *mBdnf* therapy promotes neurogenesis and the retention of hippocampal neurons without impacting amyloid plaque development.

**Figure 5:**
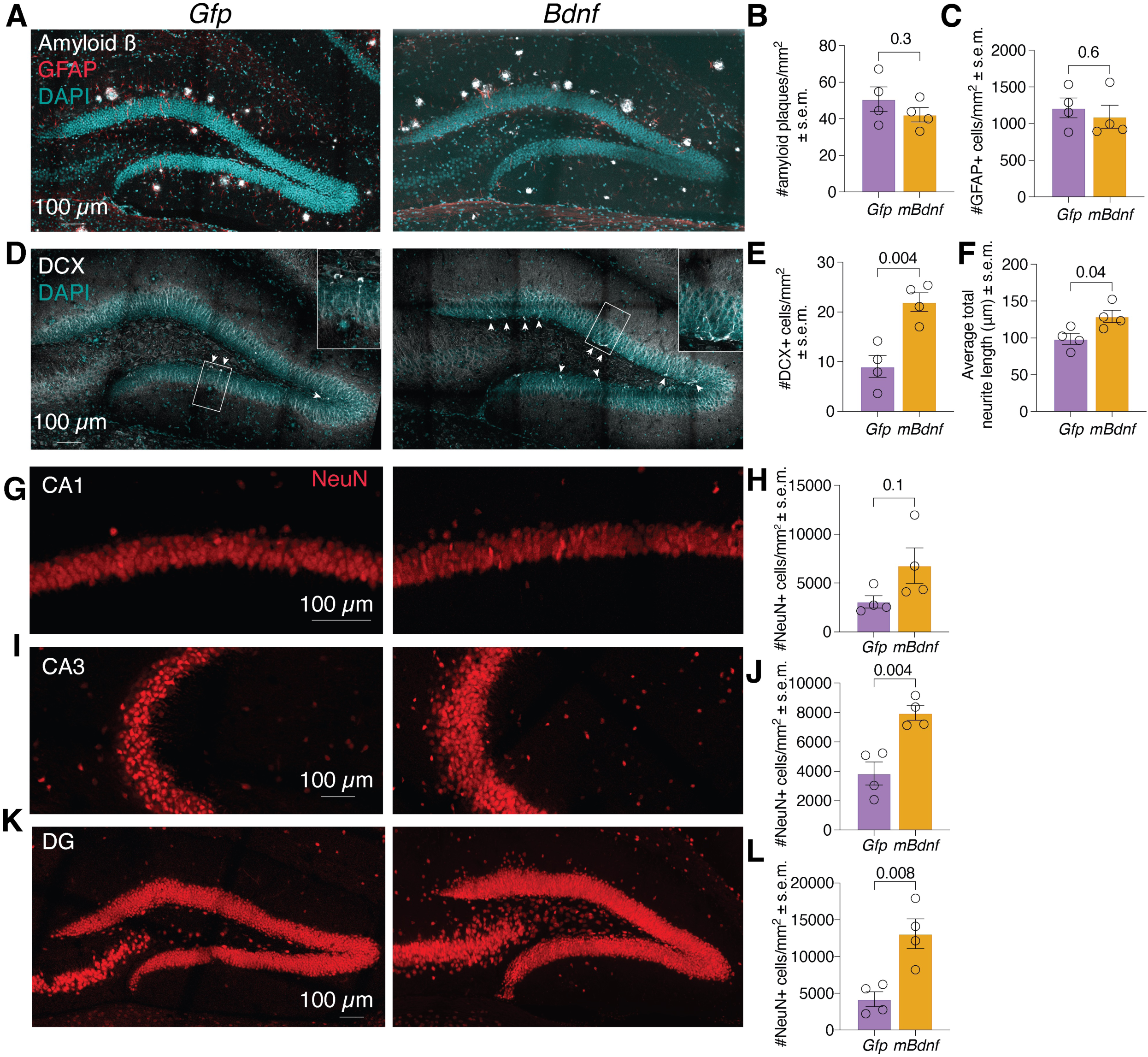
Histological analysis of *mBdnf* mRNA-LNP treatment in 5xFAD mice. **A-C**. Amyloid plaque (white) GFAP (red) and DAPI (cyan) staining of the hippocampus of *Gfp* and *mBdnf*-treated 5xFAD animals (A). Number of amyloid plaques (B) and GFAP+ cells (C) per mm^2^. **D-F.** DCX (white) and DAPI (cyan) staining of the hippocampus of *Gfp* and *mBdnf*-treated 5xFAD animals. DCX+ neurons indicated with arrows (H). Number (E) and average neurite total length (F) of DCX+ cells. **G-L.** NeuN staining in CA1 (G), CA3 (I) and Dentate Gyrus (DG) (K) regions of the hippocampus of *Gfp* and *mBdnf*-treated 5xFAD animals (J). Number of NeuN+ cells per mm^2^ quantified in H, J and L. Each circle in B, C,E, F, H, J represents hippocampal sections from an individual animal. Data analysed using an unpaired t-test.

## DISCUSSION

In this study, we show that intranasal delivery of LNP-encapsulated *Bdnf* mRNA enhances memory and promotes neurogenesis in aged mice and in the 5xFAD model of Alzheimer’s disease. This work aligns with extensive evidence in the literature highlighting the therapeutic potential of BDNF in the CNS^[24]^. Several recent studies have shown that intranasally delivered BDNF protein can improve neuronal function across multiple disease models; intranasally BDNF was shown to reduce trigeminal hyperalgesia in a migraine model^[25]^, although only at very high protein doses (40-80 µg/kg); BDNF-loaded exosomes promoted remyelination in a model of multiple sclerosis^[26]^; BDNF-loaded extracellular vesicles improved post-ischemic recovery and synaptic plasticity^[27]^ and intranasally delivered BDNF protein reversed depression-like behaviours, enhanced hippocampal neurogenesis and reduced oxidative stress in mice^[28]^.

Despite these promising outcomes, delivery of BDNF protein has several key limitations. Nasally applied proteins are subject to rapid degradation in the mucosa, diffuse inefficiently through olfactory and trigeminal pathways and exhibit short half-lives due to receptor-mediated internalisation and degradation^[29]^. In addition, vesicle-based delivery systems impose fixed cargo limits and require cell culture-dependent manufacturing, introducing batch variability and limiting scalability (Reviewed in^[30]^).

In contrast, mRNA enables local, in situ synthesis of peptides following cellular uptake. Moreover, we have shown here mRNA-driven BDNF protein expression is detectable for up to 4 days, demonstrating prolonged therapeutic exposure from a single dose. Importantly, mRNA translation amplifies the delivered mRNA cargo, generating greater and sustained amounts of peptide compared to what can be feasibly delivered by a bolus of purified protein. Lastly, mRNA-LNPs are cost-efficient, GMP-standardised, chemically defined and highly scalable, making them well suited for clinical translation.

It is of note that this study delivered mRNA encoding a modified *Bdnf* mRNA sequence. Endogenous BDNF is synthesised as pre-pro-BDNF and subsequently cleaved into pro-BDNF or mature BDNF (mBDNF). These isoforms exert opposing biological effects: while mBDNF activates TrkB signalling to promote synaptic plasticity and neuronal survival, pro-BDNF engages p75^NTR^, triggering apoptotic pathways, synaptic pruning and long-term depression. Consistently, elevated pro-BDNF levels are associated with learning and memory impairments^[13]^, and individuals carrying a Val66Met polymorphism, which prevents pro-BDNF cleavage into mBDNF, show similar cognitive deficits ^[31]^. The modified BDNF design used here avoids these detrimental effects. This design has also previously been shown to yield functional BDNF capable of supporting neuronal differentiation and survival ^[32]^ and here we similarly observed enhanced NSPC proliferation and neuronal differentiation in *mBdnf* mRNA-treated cells. Notably, other studies have delivered the full-length BDNF sequence, including the pro-domain, using MSC-derived vesicles^[33]^, clathrin nanoparticles^[34]^ or AAV vectors^[35]^ and reported functional recovery and cognitive improvements in models of stroke, HIV/neuroAIDS, neurodevelopmental pathology and depression. As such, intranasal delivery of full-length *Bdnf* mRNA may also have positive effects on neuronal viability, neurogenesis and memory-based behaviours.

While this work focused on BDNF, delivery of mRNA encoding other neurotrophic factors such as NGF or GDNF would also be of interest in future work. In rodent and non-human primate models of Parkinson’s, direct brain infusion of GDNF has been shown to protect dopaminergic neurons, increase dopamine levels and improve motor outcomes^[36]^. Similarly, targeted delivery of NGF to basal forebrain cholinergic neurons has been found to enhance neuronal function and prevent degeneration in animal models of AD^[37]^. As such, GDNF and NGF represent additional candidates that could be used with the LNP-mRNA platform presented here.

In aged mice, this study administered a single LNP-mRNA dose 24 hr before a behavioural test, without a pre-treatment phase. It is possible that repeated or earlier dosing would yield greater behavioural improvements. Younger animals, without baseline cognitive impairment, may also benefit from *mBdnf* mRNA-LNP therapy, although, as these animals show robust behaviour in learning and memory-based tasks, the ability to detect an effect in this age group would require more challenging tasks.

In 5xFAD mice we found that *mBdnf* mRNA-LNP, delivered from 8 weeks of age, improved memory and increased hippocampal neuronal numbers and neurogenesis, however, it did not alter amyloid plaque burden. This aligns with previous studies showing that BDNF does not directly modify amyloid pathology^[24c]^. Instead, BDNF appears to enhance neuronal function and resilience despite ongoing plaque accumulation. As BDNF does not restore neurons already lost, therapeutic efficacy is likely to diminish in late-stage disease. Combination treatments of *mBdnf* mRNA-LNP and current first line AD therapies such as anti-amyloid antibodies may act synergistically by reducing plaque development while preserving neuronal health.

Lastly, optimisation of both the mRNA and LNP components could enhance therapeutic outcomes. mRNA modifications such as codon optimisation and reduction of secondary structure can increase mRNA yield and prolong expression ^[38]^. 5’ and 3’ UTR engineering can also influence mRNA stability and translation ^[39]^ and promote expression in specific cell types ^[40]^. LNP modifications such as ApoE-binding ligands and alternative lipid compositions could further refine cellular uptake and distribution. Nonetheless, the mRNA-LNP formulation used in this study permits efficient delivery to the olfactory and respiratory epithelium and produces substantial cognitive improvements in both aged animals and an AD mouse model. While future refinements may further enhance this system, the current formulation already exhibits surprisingly robust effects.

Taken together, these findings establish intranasal LNP-mRNA delivery as a versatile and non-invasive platform with broad therapeutic potential. Beyond BDNF, this approach may be applicable to diverse CNS diseases, offering a promising pathway toward future neuro-regenerative therapies.

## Methods

### mRNA template cloning and *in vitro* transcription

Templates encoding the coding sequences of mRNA therapeutics were purchased as gBlocks (IDT, Integrated DNA Technologies). Sequences were restriction enzyme cloned into a T7-driven mRNA expression plasmid, based on the Pfizer-BioNTech vaccine backbone and assembled plasmid sequences confirmed by Sanger sequencing. Verified plasmids were linearised by BspQ1 restriction enzyme (NEB, #R0712L) digestion and mRNA synthesised by *in vitro* transcription using the HiScribe® T7 High Yield RNA Synthesis Kit (NEB, E2080L). mRNA for *in vitro* use was generated using standard UTP. mRNA for *in vivo* use was generated using N^1^-Methylpseudouridine-5’-Triphosphate (TriLink BioTechnologies, N-1081). Any residual DNA template was removed using DNase I (37°C, 15 min). mRNA was precipitated using LiCl (NEB, E2040S) and mRNA quantity and quality assessed using a Nanodrop.

### Lipid Nanoparticle (LNP) encapsulation

*In vitro* transcribed mRNA was encapsulated into LNPs for delivery to mice. LNP-mRNA was made up to the following composition: 47.5% ALC-0315, 10% DSPC Helper lipid, 40.5% Cholesterol, 1.8% ALC-0159 (PEG) and 0.2% DiR dye in a microfluidic mixer (NanoAssemblr^TM^ Ignite^TM^ nanoparticle formulation system, Cytiva) and at a target molar ratio of 6:1 (cationic lipid:mRNA). The LNP-mRNA solution was exchanged for 10x volume 0.01M Tris buffer using Amicon® Ultra-15 Centrifugal filters (Millipore, UFC903008) and centrifuged (2000*g*) to remove ethanol. Processed LNP-mRNA complexes were stored in 0.01M Tris and 10% (w/vol) sucrose. Encapsulation efficiencies were assessed using the Quant-iT^TM^ RiboGreen RNA assay (Invitrogen, R11490). The size, uniformity and zeta potential of LNPs was determined using a Zetasizer (Malvern Pananalytical).

### mRNA transfection

HEK293T or SH-SY5Y cells were transfected with mRNA using Lipofectamine^TM^ MessengerMax^TM^ Transfection Reagent (Invitrogen, LMRNA001), according to the manufacturer’s instructions.

### ELISA assay

BDNF protein levels in cell culture media supernatants were quantified using the Total BDNF Quantikine ELISA kit (In Vitro Technologies, DBNT00), according to the manufacturer’s instructions.

Anti-PEG antibody levels were assessed in the serum of treated animals using the Mouse anti-PEG IgG ELISA kit (Creative Diagnostics, DEIA6159), according to the manufacturer’s instructions.

### RTqPCR

Total RNA was extracted from the olfactory bulb, respiratory epithelium and olfactory epithelium of LNP-mRNA treated mice using the Isolate II RNA Mini Kit (Bioline, BIO-52073), following the manufacturer’s instructions. RNA (100 ng) was used as a template for cDNA synthesis using the iScribe cDNA synthesis kit (Biorad, #1708891) according to the manufacturer’s instructions. RTqPCR was performed in a 384-well plate using a QuantStudio^TM^ 6 Real-Time PCR machine (ThermoFisher) using the following primers:

*Bdnf* F 5’ CATGAAGGCGCACTCCGAC 3’

*Bdnf* R 5’ GCCTTTGGATACCGGGACTTT 3’

*Gapdh* F 5’ TTCACCACCATGGAGAAGGC 3’

*Gapdh* R 5’ CCCTTTTGGCTCCACCCT 3’

Samples were run using the following cycling conditions: Initial hold (50°C, 2 min), 40x cycles (95°C, 15 sec; 60°C, 1 min; 72°C, 1 min) and melt curve analysis (72°C, 2 min; 95°C 1 min; 60°C 1 min). Data was analysed using the 2^−ΔΔCT^ method ^[41]^.

### Western Immunoblotting

Freshly dissected brain tissue from *mBdnf* or *Gfp* LNP-mRNA-treated animals was dissociated into a single cell suspension using a tissue homogeniser in RIPA buffer (1% (vol/vol) Triton-X 100, 0.1% (w.vol) SDS, 1% (w/vol) sodium deoxycholate, 10 mM Tris-HCl pH 7.5, made up in H_2_O) contained 1x cOmplete^TM^ EDTA-free protease inhibitors (Merck, 1187358001). Samples were incubated at 4°C for 10 min on a roller and centrifuged (10,000g, 10 min) to remove cellular debris. The supernatant was transferred to fresh tubes. Total protein concentrations were determined using a Pierce^TM^ BCA Protein Assay kit (ThermoFisher, #23225) according to the manufacturer’s instructions. A total of 20 μg protein per sample was loaded onto a 4-20% Tris-Glycine SDS-PAGE gel (Bio-Rad, #4561094), electrophoresed and transferred onto a nitrocellulose membrane (LI-COR, 926-31092). Membranes were blocked in TBS-based Intercept® Blocking Buffer (LICOR, NC1660550) and incubated O/N at 4°C with primary antibodies against BDNF (Invitrogen, PA5-85730; 1:500 and alpha tubulin (Abcam,ab7291 1:1000). Following TBST washes, membranes were incubated for 1 hr at RT IRDye® 800CW Goat anti-rabbit IgG and IRDye® 680RD Goat anti-mouse IgG at 1:10,000. Membranes were washes, imaged using the Odyssey® Infrared Imaging System (LI-COR) and analysed using Image Studio software (LI-COR)

### Neural Stem and Progenitor Cell (NSPCs) isolation and culture

Hippocampi from 12-month-old C57Bl/6 mice were isolated under a dissecting microscope. Tissue was incubated with Accutase^TM^ (Stemcell, #07922) for 30 min on ice, followed by 10 min at 37°C and mechanically dissociated using a pipette. Dissociated tissue was passed through a 70 μm cell strainer (Corning, 431751) into cell culture plates containing NeuroCult^TM^ proliferation media (Stemcell, #05702), supplemented with 800 ng/ml Heparin (Stemcell, #07980), 20ng/ml EGF (Stemcell, #78006.1) and 20 ng/ml FGF (Stemcell, #78003.1). Cells were grown as free-floating neurospheres in 37°C and 5% CO_2_. Cells were passaged every 5-7 days using Accutase^TM^ and replated at 10,000 cells/cm^2^.

### SH-SY5Y culture

SH-SY5Y cells were grown at 37°C in 5% CO_2_ in DMEM/F12 (Gibco, 11320033) supplemented with 1x GlutaMAX^TM^ (35050061), 1% (v/v) penicillin-streptomycin (Gibco, 15140122) and 10% (v/v) FBS. Cells were passaged using TrypLE and replated at 10,000 cells/cm^2^.

### SH-SY5Y differentiation

SH-SY5Y cells were differentiated as described ^[42]^, with minor modifications. Briefly, cells were seeded at 50% confluency onto poly-D-lysine (Gibco, A3890401) and laminin (Sigma, L2020) coated chamber slides (Merck, Z734756) in DMEM/F12 (Gibco, 11320033) supplemented with 1x GlutaMAX^TM^ (35050061), 1% (v/v) penicillin-streptomycin (Gibco, 15140122) and 10% (v/v) FBS. The following morning media was replaced with conditioned media from HEK293T cells transfected with *mBdnf* or *Gfp* mRNA in Stage I media (DMEM/F12, 1x GlutaMAX^TM^, 1% (v/v) penicillin-streptomycin, 2.5% (v/v) FBS and 10 μM retinoic acid. Cells were cultured for 5 days before media was replaced with Stage II conditioned media (Neurobasal-A (Gibco, 10888022), 20 mM KCl, 1x B27^TM^ (Gibco,17504044), 1x GlutaMAX^TM^, and 1% (v/v) penicillin-streptomycin with or without 50 ng/ml recombinant BDNF (Gibco, 450-02)). Cells were cultured for an additional 5 days before downstream staining and analysis.

### Histology and Immunofluorescence

For histological analysis, mice were perfusion fixed with PBS followed by 4% (w/v) PFA (Sigma, P6148). Brains were dissected and post-fixed for 24 hr in 4% PFA (w/v) at 4°C on a roller. Brain tissue was mounted in 2% (w/v) low gelling temperature agarose (Merck, A9414) and stored at 4°C until sectioning using a Vibratome (Leica). 30 μm coronal sections were blocked and permeabilised in 0.03% (vol/vol) Triton-X 100 and 10% (vol/vol) FCS for 1 hr at RT and incubated O/N at 4°C in 10% FCS containing primary antibodies. The following morning sections were washed in PBS (3x 5 min washes) and incubated with species-specific secondary antibodies for 1 hr at RT. Sections were washed, incubated with DAPI for 10 min at RT, washed and mounted onto Superfrost^TM^ slides (Bio-Strategy Pty Limited, EPBRSF41296SP) in ProLong^TM^ Diamond antifade mounting medium (Invitrogen, P36962).

| Antibody | Source | Catalogue # | Dilution |
| --- | --- | --- | --- |
| Anti-NeuN A488 | Abcam | Ab190195 | 1:1000 |
| Anti-Amyloid beta | Cell Signalling | 2450T | 1:1000 |
| Anti-GFAP | Abcam | Ab68428 | 1:500 |
| Anti-DCX | Abcam | Ab18723 | 1:200 |
| Anti-BDNF | Abcam | Ab108319 | 1:500 |
| Goat anti-mouse<br>A488 | Invitrogen | A110011 | 1:400 |
| Goat anti-rabbit<br>A488 | Invitrogen | A21428 | 1:400 |
| Goat anti-rabbit<br>A647 | Invitrogen | A21245 | 1:400 |

### Image acquisition

Confocal images were acquired using a Marianas 3i spinning disk confocal microscope. 20-40 μm Z-stacks were acquired at 0.34 μm (40x magnification) or 0.78 μm (20x magnification) steps. Maximum Intensity Projections were generated and analysed in Image J (Version 2.16.0/1.54j). Cell numbers were quantified on auto-thresholded images using the ‘Analyze Particles’ function. DCX neurite analysis was performed using the NeuronJ plugin.

### Mouse Ethics

This study was conducted in accordance with the guidelines set out in the Australian Code of Practise for the Care and Use of Animals for Scientific Purposes. All experiments were conducted with approval of The University of Sydney Animal Ethics Committee. Mice were housed 2-5 to a cage with standard cage enrichments and provided with food and water ad libitum. Mice were housed in ventilated cages on a 12-hr light/dark cycle.

### Intranasal administration

Mice were lightly anaesthetised using 2.5% isoflurane and 1% O_2_. Mice were gently scruffed and 5 μl mRNA-LNP, containing 0.5 μg mRNA was administered dropwise to each nostril using a P20 pipette.

### Mouse behavioural tests

For behavioural testing, light levels were standardised at 100 lux using a lumometer (Lutron LM-81LX, S041136) and the temperature maintained at 22°C. Behavioural tests were performed in the light phase of the light-dark cycle and only one test was performed per day. Prior to testing all mice were habituated to the operator and habituated to the room for at least 30 min prior to testing. All mice were returned to their home cage following testing. Animals were recorded from above using a webcam (WebCam C615 (Logitech; Lausanne, Switzerland) and data analysed using Any-maze software (ANY-maze, Version 7.20).

### Open field test

The open field was performed as described ^[43]^. Each mouse was placed into a 40cm^2^ square arena with 30cm high black walls. Mice were allowed to freely explore for 10 min and ambulation assessed and the proportion of time spent at the periphery of the arena was used as a measurement of anxiety.

### Novel Object recognition

The novel object recognition test was performed as described ^[44]^. Briefly, each mouse was placed into a 40cm2 square arena with 30cm high black walls. In session 1 animals were given two identical objects and allowed to freely explore for 10 min. Animals were returned to their home cage for 2 hr. Upon returning to the arena, one of the objects from session 1 was replaced with a novel object in session 2. Animals were allowed to freely explore for 10 min. The ratio of time spent with the novel versus familiar object in session 2 was used to assess spatial memory.

### Y maze for working memory and spatial memory

Two tests were formed in the Y maze, as described ^[45]^. In both tests each mouse was placed into a Y shaped maze with three arms at 120° to each other. Each arm was 30cm long, 5 cm wide and 20 cm high. Visual cures were placed at the end of each arm to assist with spatial orientation. Animals began both tests in the ‘home arm’. The other two arms were defined as arm A and B. In the Y maze for working memory animals were allowed to freely explore the maze for 5 min. The sequence of entries into each arm was assessed, with one ‘spontaneous alternation’ defined as a mouse having entered each of the three arms within three consecutive arms entries. The proportion of spontaneous alternations was defined as the number of alternations/(total number of arm entries – 2) and used as a measurement of working memory.

In the Y maze for spatial memory, in session 1, each mouse was placed into the Y maze with either arm A or B blocked from entry (novel arm). Animals were allowed to freely explore the remaining two arms for 10 min. Animals were returned to their home cage for 2 hr. Upon return to the Y maze all arms were open, and animals were allowed to freely explore for 5 min. The time spend in the previously closed ‘novel arm’ compared to the familiar arm, was used as a readout of spatial memory.

## Author Contributions

MB and GN designed the study.

RMDH provided materials and experimental input.

MB, TC, LL, KF, RC, HS and AP performed experiments and analysed data.

GN supervised the work.

All authors contributed to the manuscript.

## Acknowledgements

The authors would like to acknowledge the Sydney imaging facility for technical assistance and ABR and the Charles Perkins Center animal technicians for excellent animal care.

