## Supplemental Data for "Non-invasive *Bdnf* mRNA therapy improves cognition in ageing and Alzheimer’s mouse models"

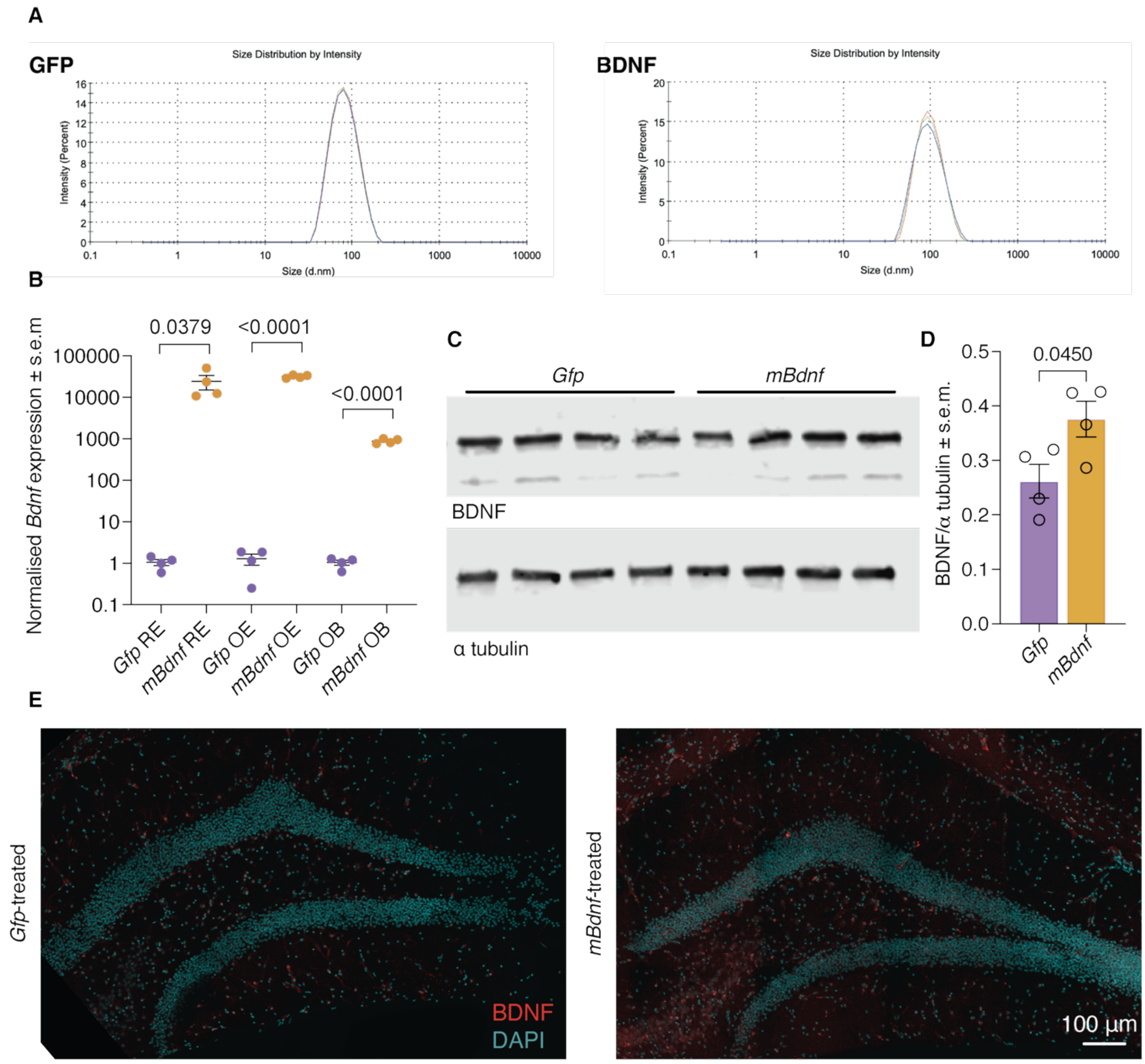


**Fig S1**

**(A)** Average diameter (nm) of LNPs encapsulating mRNA encoding GFP or BDNF. Size determined using a zetasizer.

**(B)** *Bdnf* mRNA expression in the RE (respiratory epithelium), OE (olfactory epithelium) and OB (olfactory bulb) of animals treated with LNP-mRNA encoding *Gfp* of *mBdnf*. Tissue collected 24 hr after LNP-mRNA intranasal administration. *Bdnf* levels normalised to *Gapdh* housekeeping control and data normalised to *Gfp*-treated controls. Each circle represents cDNA isolated from an individual animal. Data analysed using unpaired t-tests.

**(C,D)** Western Immunoblot of BDNF and $\alpha$-tubulin (loading control) in total protein lysates of brains from *Gfp* or *mBdnf*-treated animals (C). Densitometry analysis is shown in (D). Animals treated with 1 $\mu$g mRNA-LNP. Tissue collected 24 hr after treatment.

**(E)** BDNF (red) and DAPI (cyan) staining in hippocampal sections from *Gfp* or *mBdnf*-treated mice. Representative images of 3-4 animals per treatment group.

Each circle in D represents one lane of the corresponding western immunoblot in C. Data analysed using an unpaired t-test.

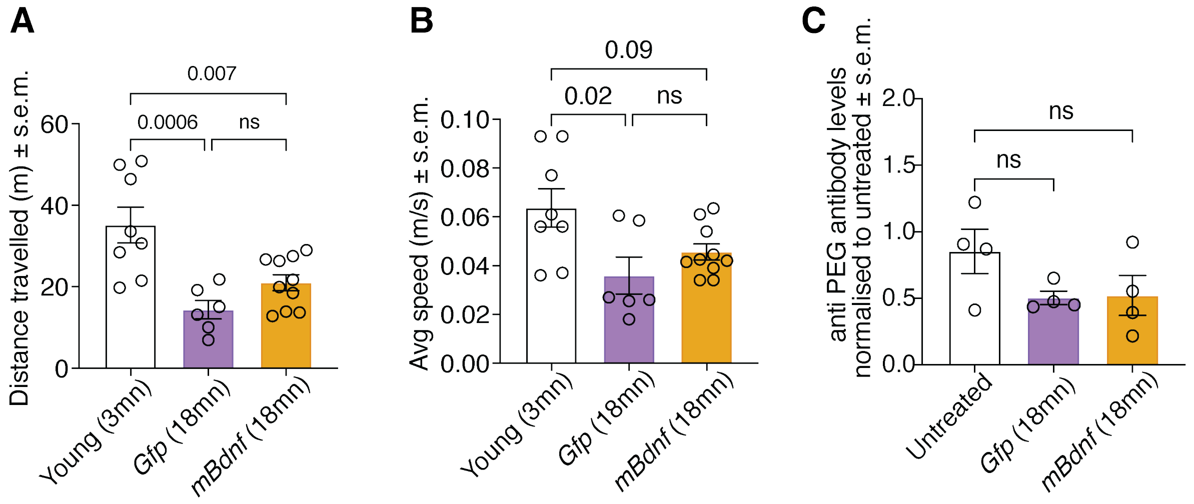


**Fig S2**

**(A)** Distance travelled (m) by young (3mn) mice or aged (18mn) mice treated with *Gfp* or *mBdnf* LNP-mRNA in the open field test.

**(B)** Average speed (m/s) of young (3mn) mice or aged (18mn) mice treated with *Gfp* or *mBdnf* in the open field test.

**(C)** Normalised levels of anti-PEG antibodies in the serum of aged (18mn) animals treated with *Gfp* or *mBdnf* and untreated controls. Serum collected after 4x LNP-mRNA administrations.

Each circle represents an individual animal. Data analysed using a one-way ANOVA with Dunnett post-hoc correction.

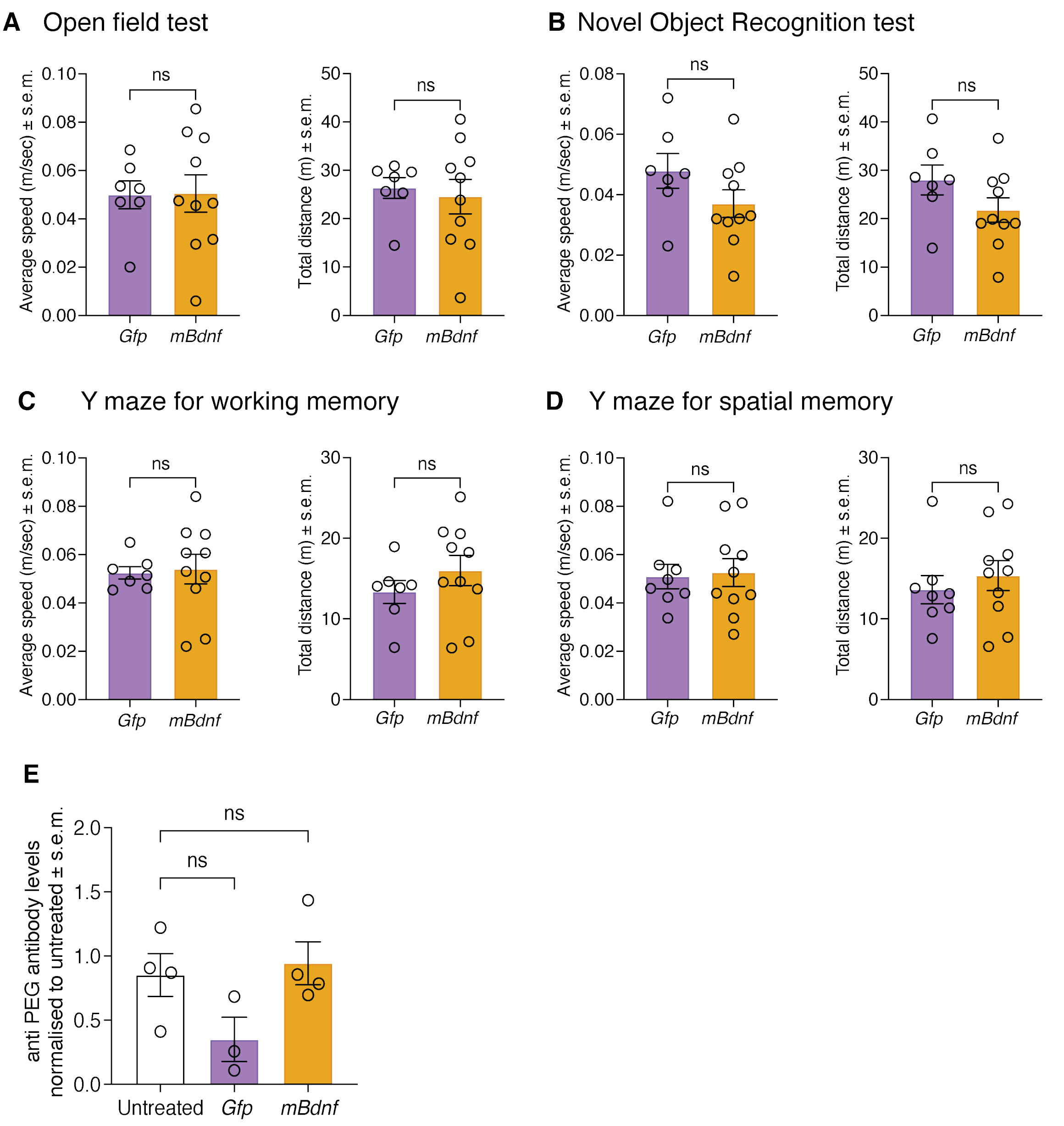


**Fig S3**

**(A-D)** Average speed (m/s) and distance (m) measurements for *Gfp* and *mBdnf*-treated 5xFAD animals in the open field test (A), novel object recognition test (B), Y maze for working memory (C) and Y maze for spatial memory (D).

**(E)** Normalised levels of anti-PEG antibodies in the serum of 5xFAD animals treated with *Gfp* or *mBdnf* and untreated controls. Serum collected after 20x LNP-mRNA administrations.

Each circle represents an individual animal. Data analysed using unpaired t tests (A-D) or a one-way ANOVA with Dunnett post hoc correction (E).
